# Behavioural cue reactivity to alcohol and non-alcohol-related stimuli among individuals with severe alcohol use disorder: an fMRI study with a visual task

**DOI:** 10.1101/2020.02.03.932889

**Authors:** Shou Fukushima, Hironori Kuga, Naoya Oribe, Takeo Mutou, Takefumi Yuzuriha, Hiroki Ozawa, Takefumi Ueno

**Affiliations:** Department of Neuropsychiatry, Nagasaki University, Nagasaki, Japan; Department of Clinical Research, National Hospital Organization, Hizen Psychiatric Medical Center, Saga, Japan; Department of psychiatry, Michinoo Hospital, Nagasaki, Japan; Department of Neuropsychiatry, Kyushu University, Fukuoka, Japan

## Abstract

Patients with alcohol use disorder (AUD) have difficulties controlling alcohol cravings and thus exhibit increased use and earlier relapse. Although patients tend to respond more strongly to alcohol-related images compared with non-alcohol-related images, few researchers examined the factors that modulate cravings. Here, we examined whole-brain blood oxygen level-dependent (BOLD) responses to behavioural cues in individuals with AUD and healthy controls (HCs). The participants included 24 patients with severe AUD and 15 HCs. We presented four beverage images (juice, drinking juice, sake, and drinking sake) and compared participant BOLD responses between the two groups. Multiple comparisons revealed that the AUD group had lower BOLD responses compared with the HC group to images of drinking juice in the left precuneus (p = 0.036) and the left posterior cingulate cortex (PCC) (p = 0.044) and higher BOLD responses to images of drinking sake in the left PCC (p = 0.044). Furthermore, compared to the HCs, the AUD patients had decreased BOLD responses associated with cue reactivity to drinking juice in the left precuneus during the period from 15 to 18 s (p = 0.004, df = 37) and 18 to 21 s (p = 0.002, df = 37). Using the Spearman correlation, we found a significant negative correlation between BOLD responses in the left PCC of the AUD patients and Mini–Mental State Examination (MMSE) scores (r = −0.619, p = 0.001). Our findings suggest that HCs and severe AUD patients differ in their responses not to images of alcoholic beverages but those related to alcohol drinking behavior. Thus, these patients appear to have different patterns of brain activity. This information may aid clinicians in developing treatments for patients with AUD.

## Introduction

According to the World Health Organization (WHO), 3.3 million deaths every year result from the harmful use of alcohol, which represents 5.9% of all deaths in the world [1]. The harmful use of alcohol is a causal factor in more than 200 diseases and injuries [2]. Overall, 5.1% of the global burden of disease and injury is attributable to alcohol, as measured in disability-adjusted life years [1]. Beyond health problems, the harmful use of alcohol is associated with significant social and economic losses for individuals and society at large.

As the American Psychiatric Association (APA) guidelines suggest, not only abstinence from alcohol use but also reduction or moderation of alcohol use may be appropriate initial goals of treatment for alcohol use disorder (AUD) from a harm reduction perspective. In substance use research, however, exposure to substance-related cues (such as the sight or smell of alcoholic beverages) has been found to evoke elevations in subjective craving and physiological arousal and increase the likelihood of substance use [3]. Thus, craving appears to play an important role in predicting negative outcomes such as increased use and earlier relapse in AUD patients [4]. However, some patients are not sensitive to the craving before their relapse. Furthermore, the mechanisms underlying craving remain unclear.

Previous studies investigating craving have suggested that the posterior cingulate cortex (PCC) plays a role in craving and relapse to alcohol use [5,6]. The PCC is a primary node of the default mode network (DMN) [7,8], which might be related to deficits in brain function in AUD patients. The PCC codes relevant information from visual sensory systems to evaluate emotional content [9] and is involved in internally directed cognition, such as memory retrieval and planning [10]. This region plays a crucial role in integrating incoming episodic memory with existing knowledge to create a coherent representation of the event [11]. While the PCC is a well-studied region with AUD patients in task-based studies on craving, and AUD patients have been found to have greater responsiveness to alcohol-related images than non-alcohol beverage images [12], few studies have examined the factors that contribute to craving in AUD patients.

Given the findings of several reports using human brain imaging, we hypothesized that the presentation of stimuli involving alcohol drinking behavior may be accompanied by stronger brain activation in AUD patients compared with stimuli that simply contain alcoholic beverages. To address the question of whether and how the PCC and other brain regions related to the DMN are implicated in the deficits, we examined possible alterations in brain function in patients with AUD using functional MRI (fMRI) with beverage image cues. We anticipated that such alterations might be correlated with changes in cognitive function in the patient group.

## Materials and methods

### Participants

Demographic and clinical data are shown in Table 1. The sample consisted of 15 healthy controls (HCs) and 23 patients with severe AUD and one patient with moderate AUD. All participants were between 25 and 60 years of age. The HCs were screened using the Structured Clinical Interview (SCID)–non-patient edition [13].

**Table 1.** Demographic and Clinical Characteristics of the Participant Groups.

|  | AUD (N = 24) | HC (N = 15) | F or t | df | p-Value |
| --- | --- | --- | --- | --- | --- |
| Age (mean, SD) | 47.5 ± 8.6 | 46.7 ± 7.9 | 0.30 | 37 | 0.77 |
| Sex (male), N (%) | 17 (71) | 10 (67) | 0.27 | 37 | 0.79 |
| Handedness<sup>a</sup> (mean, SD) | 87.7 ± 39.5 | 89.5 ± 14.9 | 1.80 | 37 | 0.87 |
| Years of education (mean, SD) | 12.8 ± 2.4 | 16.4 ± 2.9 | 0.00 | 37 | < 0.001★ |
| AUDIT score (mean, SD) | 28.3 ± 7.1 | 1.7 ± 2.5 | / | / | < 0.001 |
| DSM-5 criteria met (mean, SD) | 8.9 ± 2.1 | 0.20 ± 0.6 | / | / | < 0.001 |
| MMSE score (mean, SD) | 28.5 ± 1.6 | / |  |  |  |
| Duration of illness (mean, SD) | 8.7 ± 6.9 | / |  |  |  |
| Number of hospital admissions (mean, SD) | 2.5 ± 1.4 | / |  |  |  |
| Neurologic symptoms<sup>b</sup> (yes), N (%) | 10 (42) | / |  |  |  |

**Note: Values are means ± SD unless otherwise indicated. HC, healthy control; AUD, alcohol use disorder; AUDIT, Alcohol Use Disorders Identification Test; MMSE, Mini-Mental State Examination.**
<sup>a</sup>Based on the Annett Handedness Scale total score.
<sup>b</sup>Past history of withdrawal convulsions or hallucinations.
★Patients with AUD had significantly fewer years of education compared to HCs via a t-test.

**Seven patients were medicated with antipsychotics (only atypical antipsychotics; chlorpromazine equivalents = 180.5 ± 132.3 mg), and 17 patients were unmedicated. Within the patient group, three participants received acamprosate (1443 ± 508.7 mg), and six received antidepressant drugs.**

**Ten patients had a past history of neurologic symptoms, such as withdrawal convulsions or hallucinations.**

The participants with AUD were recruited from a pool of patients who had been hospitalized at our institution and who subsequently joined the alcohol rehabilitation program (ARP) to recover from AUD. All AUD patients were screened using the Alcohol Use Disorders Identification Test (AUDIT; [14). They were then diagnosed via the DSM-5 [15] criterion as having severe AUD. We measured cognitive impairment using the Mini–Mental State Examination (MMSE). We measured the MRI in AUD participants 1–2 months after admission in order not to be confounded by withdrawal convulsions or hallucinations.

### Design and ethical approval

This study had a prospective, intervention-based cross-sectional design. The study intervention involved presenting images of beverages (alcoholic and non-alcoholic). The protocol and informed consent form were approved by the ethics committee of the National Hospital Organization Hizen Psychiatric Center. After receiving a complete description of the study, all participants signed an informed consent form.

### Stimuli and procedure

The methods were based on a previous study [16]. In the informed consent process, we explained to all participants that the tasks included the presentation of images, but we did not describe the content of the images. The participants completed eye tests, and the participants with weak sight were given glasses to wear during the MRI.

The participants were asked to lie in a supine position on a bed inside the MRI scanner while wearing headphones. The head of each participant was restrained via padding behind the neck and between the head and the coil. Participants were asked to keep their head still inside the scanner and to focus on a fixation cross presented on a screen. All visual stimuli were presented on the screen.

We presented the visual stimuli using a two-min block-design paradigm with eight blocks of 15 s of rest (viewing a mosaic image) and eight blocks of 15 s of stimulus presentation (viewing a beverage image). We presented four beverage images (Image A, juice; Image B, drinking juice; Image C, sake; and Image D, drinking sake) that varied in terms of valence. We previously created a table that arranged the four beverage images into random sequences. This table was used to assign random test sequences to the participants. Sake is a type of rice wine, and patients with severe AUD in Japan often drink this kind of alcohol because it has a high alcohol content and is cheap, portable, and convenient to drink; 19 of the 24 AUD participants in this study stated that sake was their preferred alcoholic beverage (Fig 1),.

**Fig 1.**
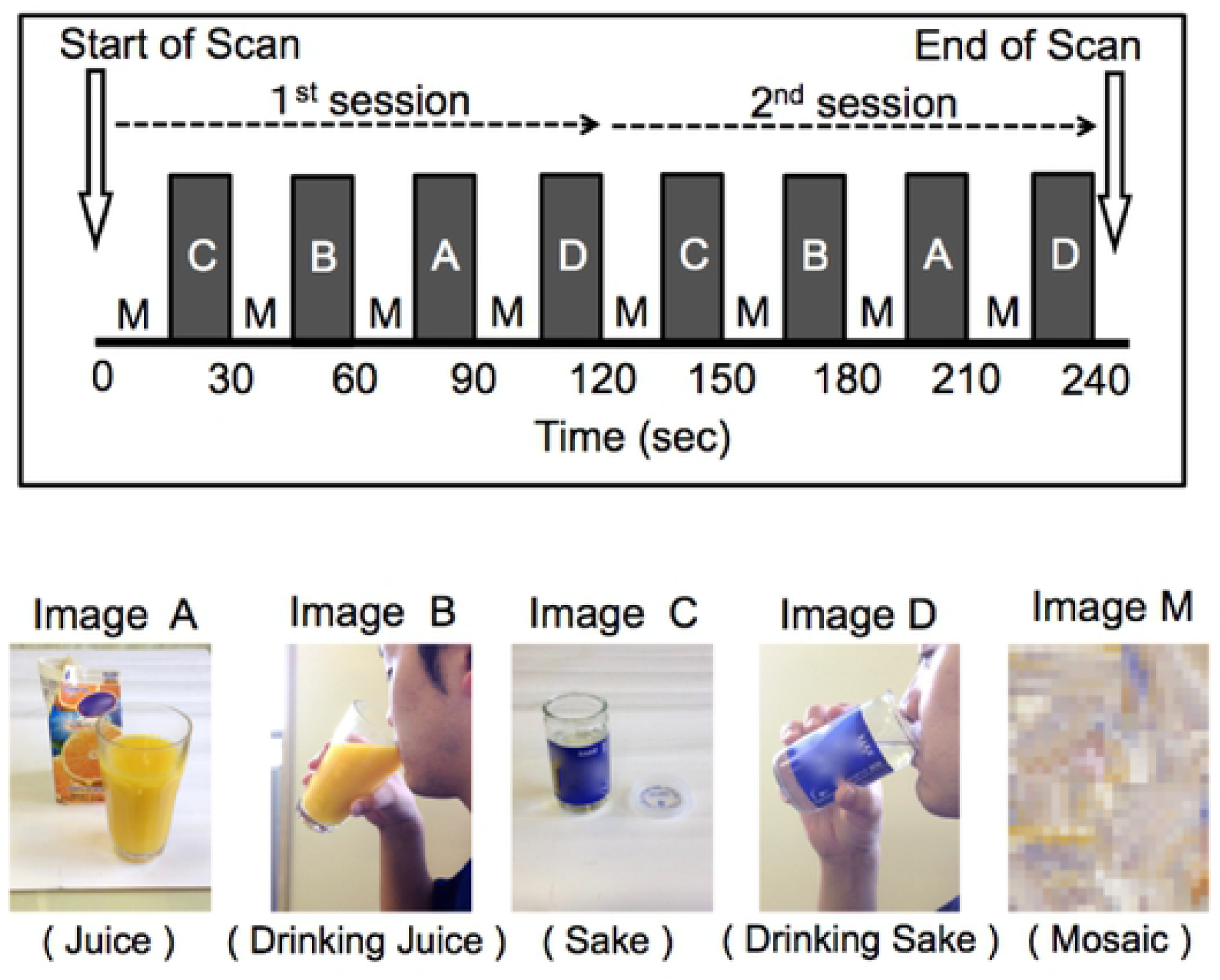
Study design and presenting images. We used a block design, and presented four beverage images (Image A, juice; Image B, drinking juice; Image C, sake; and Image D, drinking sake) and a mosaic image (Image M) to participants. Sake is rice wine. One session consisted of four rest blocks and four beverage blocks, and two sessions took 240 s. Beverage images were presented in order of valence (for example, “C-B-A-D” in this figure), and each beverage image was presented twice in each fMRI session. In each MRI session, we presented the brand of each beverage in the images with permission from the copyright holder. Due to the requests from the copyright holders of the juice and sake brands that we used, we obscured the beverage brands via a mosaic in this report.

Each block used one image. The order of images was counterbalanced across participants, and each beverage image was presented twice. In total, we presented eight stimuli for two minutes in each fMRI session.

### Data acquisition

We conducted MRI using a 1.5-T Philips scanner with a standard head coil located at the National Hospital Organization Hizen Psychiatric Medical Center. We used standard sequence parameters to obtain functional images as follows: gradient-echo echo-planar imaging (EPI); repetition time (TR) = 3000 ms; echo time (TE) = 45 ms; flip angle = 90°; field of view (FOV) = 230 × 230 mm; matrix = 64 × 64; 60 axial slices with a slice thickness of 4 mm with no slice gap. We acquired a high-resolution T1-weighted 3D anatomical image for each participant between the functional data trials.

### Image processing

Raw image DICOM files were converted to the NIFTI format using MRI-Convert (Version 2.0, Lewis Center for Neuroimaging, Oregon). Image processing and statistical analyses were performed using the statistical parametric mapping software SPM12 [17] with MATLAB R2015a. The first five volumes of each EPI image run were excluded to allow the MR signal to reach a state of equilibrium. All volumes of the functional EPI images were realigned to the first volume of each session to correct for participant motion. These images were managed with slice-timing correction, and the mean functional EPI image was then spatially co-registered with the anatomical T1 images. Each co-registered T1-weighted anatomical image was normalized to a standard T1 template image (ICBM 152), which defined the Montreal Neurological Institute (MNI) space. The parameters from this normalization process were then applied to each functional image. The normalized functional images were smoothed with a 3D 8-mm full-width half-maximum (FWHM) Gaussian kernel. Time series data at each voxel were temporally filtered using a high-pass filter with a cutoff of 128 s.

### Statistical analysis

We used one-way analyses of variance (ANOVAs), t-tests, and Wilcoxon rank-sum tests to assess group differences in the demographic variables. We performed fMRI statistical analysis on the preprocessed EPIs with the general linear model (GLM) using a two-level approach [18]. The model consisted of boxcar functions convolved with the canonical hemodynamic response function, which were then used as regressors in the regression analysis. Six head motion parameters, derived from realignment processing, were also used as regressors to reduce motion-related artifacts. On the first level of analysis, individual contrast images for each stimulus versus rest were computed and taken to the second level for random-effects inference. On the second level, contrast images for stimuli as the within-subject factors were submitted to two groups (AUD and HC) as the between-subject factors in a full-factorial ANOVA with education as a covariate. All fMRI results are reported at a significance level of p < 0.05, family-wise error (FWE)-corrected (voxel-level corrected), or p < 0.05, FWE cluster-corrected across the whole brain with the initial voxel threshold at p < 0.001.

We extracted contrast values by identifying the precuneus and the PCC as the regions of interest (ROIs) using MarsBar (http://marsbar.sourceforge.net). We chose these areas because they contained significant stimulus-by-group interactions. For each stimulus, we compared the BOLD responses in the ROIs between the two groups (AUD and HC) via SPSS (ver. 24). To test the differences between the two groups, we used t-tests for data that were normally distributed and the Wilcoxon rank-sum test for data that were not normally distributed. We applied the false discovery rate (FDR) to the results of the t-tests or Wilcoxon rank-sum tests to examine multiple comparisons using R, which is a programming language and free software environment (https://www.r-project.org/about.html). These comparison results are reported at a significance level of p < 0.05.

We used mixed-effect regression models in SPSS (ver. 24) with random intercepts to test our hypothesis that the time course of brain activities in the AUD and HC groups would differ with respect to the behavioural cue reactivity of drinking juice with education as a covariate. We chose the BOLD responses to the cue of drinking juice in the left precuneus because we found a significant time-by-group interaction in that region. To test the differences between these two groups, we used t-tests with a significance level of p < 0.05, Bonferroni corrected.

In the AUD group, we used the Spearman correlation in SPSS (ver. 24) to determine the relationship between MMSE scores and the BOLD responses associated with cue reactivity to the drinking alcohol image.

### Outcome

The primary outcome was the BOLD responses to beverage images (images of beverages or images of drinking beverages). The BOLD response was measured using the statistical parametric mapping software SPM12 (Wellcome Department of Cognitive Neurology, London, United Kingdom) with MATLAB R2015a (The Math Works Inc., Natick, MA). The secondary outcome was the correlation between MMSE scores and the BOLD responses associated with cue reactivity to the drinking alcohol image.

## Results

### Group demographic data

Means, SDs, and frequencies for clinical and demographic variables are presented in Table 1. We found no significant group differences in the demographic data with respect to age, sex, and handedness. However, the participants with AUD had significantly lower levels of education compared with the HCs (p < 0.001, F = 0.0, df = 37), as well as higher AUDIT scores and number of criteria for AUD met from the DSM-5 (p < 0.001, F =14.398, df = 37). There were no differences in the number of hospital admissions, comorbid conditions, or preferred alcohol type.

### Main effects of group and image and stimulus-by-group interactions

As shown in Table 2 and Fig 2, repeated measures ANOVAs showed several significant group differences. The main effect of group was associated with significant differences in activity in the right precentral cortex (10, −26, 62; cluster size = 2,416; F[11,28] = 25.32; cluster-level, FWE-corrected p < 0.001), the left precentral cortex (−2, −26, 66; cluster size = 2,416; F[11,28] = 14.06; cluster-level, FWE-corrected p < 0.001), (−4, −18, 58; cluster size = 712; F[11,28] = 25.32; cluster-level, FWE-corrected p < 0.001), the left supramarginal cortex (−60, −40, 24; cluster size = 2,416; F[11,28] = 15.74; cluster-level, FWE-corrected p = 0.025), the left cerebellum exterior (−8, −36, −22; cluster size = 1,008; F[11,28] = 15.05; cluster-level, FWE-corrected p < 0.001), the left hippocampus (−22, −30, −10; cluster size = 1,008; F[11,28] = 14.44; cluster-level, FWE-corrected p < 0.001), and the exterior right cerebellum (4, −36, −22; cluster size = 1,008; F[11,28] = 13.30; cluster-level, FWE-corrected p < 0.001). Furthermore, we found a significant stimulus-by-group interaction in the left precuneus, the right precuneus, and the left PCC (cluster-level uncorrected p = 0.002).

**Table 2.** fMRI Results for Anatomical Region, Seed Voxel Coordinates (MNI), and F-Values for Significant Stimulus-by-Group Interactions.

| Cluster Size (mm<sup>3</sup>) | MNI coordinates | | | $K_E$ | F-value | Anatomical Region |
| --- | --- | --- | --- | --- | --- | --- |
|  | x | y | z |  |  |  |
| 832 | -2 | -54 | 18 | 115 | 5.71 | Left precuneus |
|  | 6 | -54 | 20 |  |  | Right precuneus |
|  | 0 | -48 | 24 |  |  | Left posterior cingulate cortex |

**BOLD, blood oxygen level-dependent; MNI, Montreal Neurological Institute.**
**Results are thresholded at $p < 0.05$ , cluster-corrected.**

**Fig 2.**
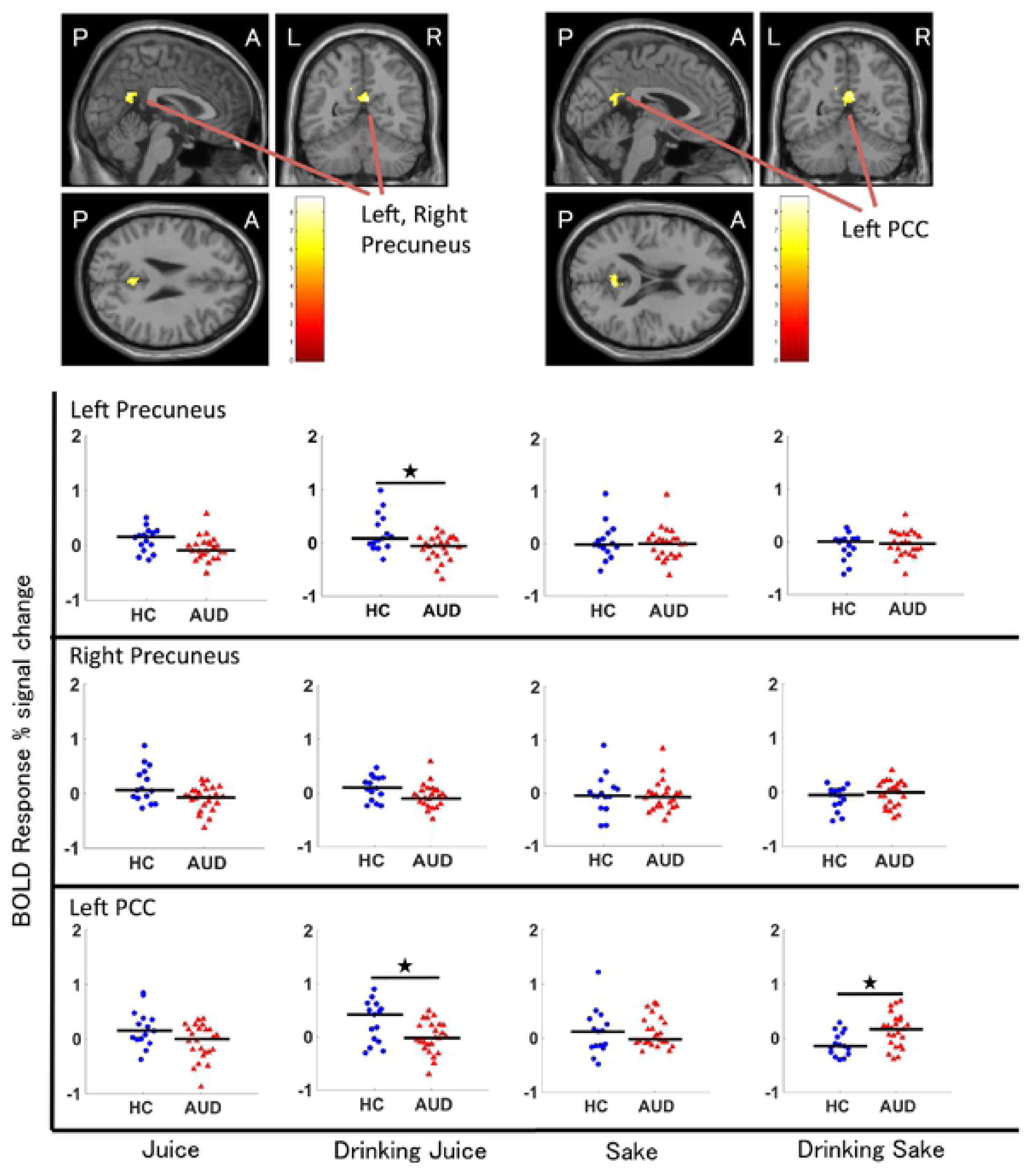
Regions where fMRI analyses revealed significant stimuli-by-group interactions. Colored bars represent the F-values of the interactions (cluster-level FWE corrected *p* < 0.05). The scatter diagram shows the BOLD responses to the four beverage images (juice, drinking juice, sake, and drinking sake) in healthy controls [(HCs); n = 15, blue dots] and participants with severe AUD [(AUD); n = 24, red dots] in the left precuneus, right precuneus, and left PCC. The bars show the median values. ^★^*p* < 0.05.

### BOLD contrast for behavioural cue reactivity associated with drinking juice vs. drinking sake in the precuneus and PCC

To determine the direction of the stimulus-by-group interaction, we extracted contrast values by identifying the PCC and precuneus as ROIs. A post hoc test with multiple comparisons revealed the following significant group differences for Image B (drinking juice) and Image D (drinking sake); HCs > AUD in terms of BOLD responses to Image B in the left precuneus (p = 0.036, t = −3.12) and left PCC (p = 0.044, t = −2.67), and AUD > HCs in terms of BOLD responses to Image D in the left PCC (p = 0.044, t = 2.73). In the right precuneus, however, our test showed no significant differences in BOLD responses between the participants with AUD and HCs. Furthermore, in the three ROIs, our tests showed no significant differences in BOLD responses to Image A (juice) and Image C (sake) between the AUD participants and HCs. Fig 2 shows the regions where the fMRI analyses revealed significant stimulus-by-group interactions and a scatter diagram. There was a group difference between the distribution trends.

In the left PCC, the median of the BOLD responses to Image B in the AUD participants was lower than that in HCs, but the median of the BOLD responses to Image D in the AUD participants was higher than that in the HCs. In the left precuneus, however, this inverse relationship was absent.

### The time course of BOLD activation of behavioural cue reactivity for drinking juice vs. drinking sake

We found significant group differences in the time course of BOLD responses with a linear mixed-effect model and a post hoc t-test (F = 2.726, p = 0.010). Specifically, we found that unlike the HC group, the AUD group exhibited a BOLD decrease in cue reactivity in the left precuneus during the period from 15 s to 18 s (p = 0.004, t = −3.044, df = 37) and from 18 s to 21 s (p = 0.002, t = −3.392, df = 37) after the onset of Image B. Conversely, we found no significant difference between the two groups in the time course for cue reactivity associated with Image D in the left precuneus, Image B in the left PCC, or Image D in the left PCC (Fig 3a-d).

**Fig 3.**
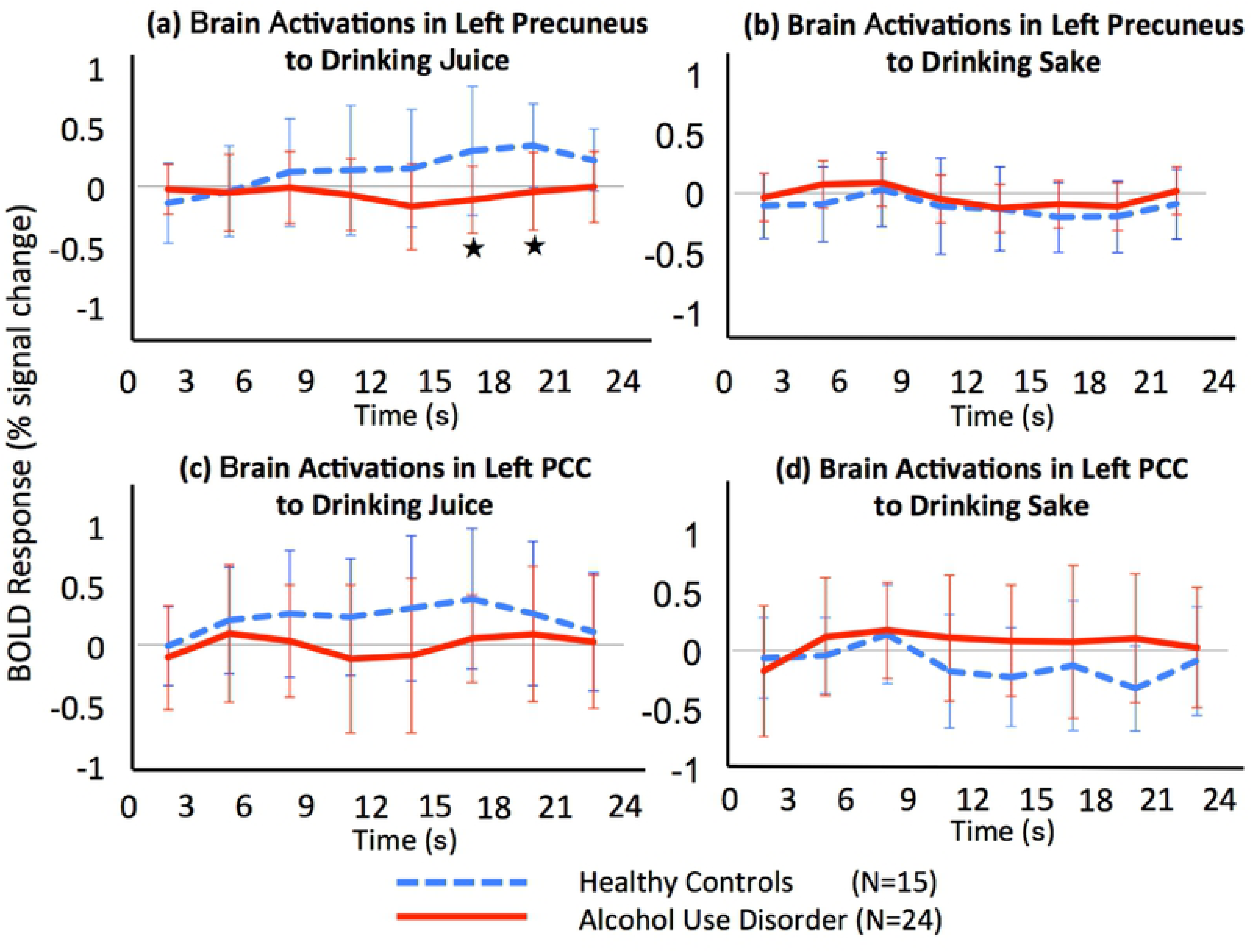
The time course of BOLD activation associated with behavioural cue reactivity for drinking juice or drinking sake in the left precuneus or left PCC. The x-axis indicates time (s), and the y-axis indicates BOLD percent signal change. The blue dotted lines and red lines indicate the BOLD percent signal change in the HCs and AUD groups, respectively. The bars show each standard error. Compared with the HCs group, the AUD group had a significantly decreased BOLD response in the left precuneus to the drinking juice stimulus from 15 to 21 s (^★^*p* < 0.0071) (a).

### Correlation between BOLD responses to the drinking sake stimulus and MMSE scores

Post hoc tests with the Spearman correlation revealed one correlation. When we conducted a correlation analysis between MMSE scores and BOLD responses in the AUD participants, we found a significant negative correlation between BOLD responses and MMSE scores in the left PCC (r = −0.619, p = 0.001) and no significant negative correlation in the left precuneus (r = −0.457, p = 0.025) (Fig 4).

**Fig 4.**
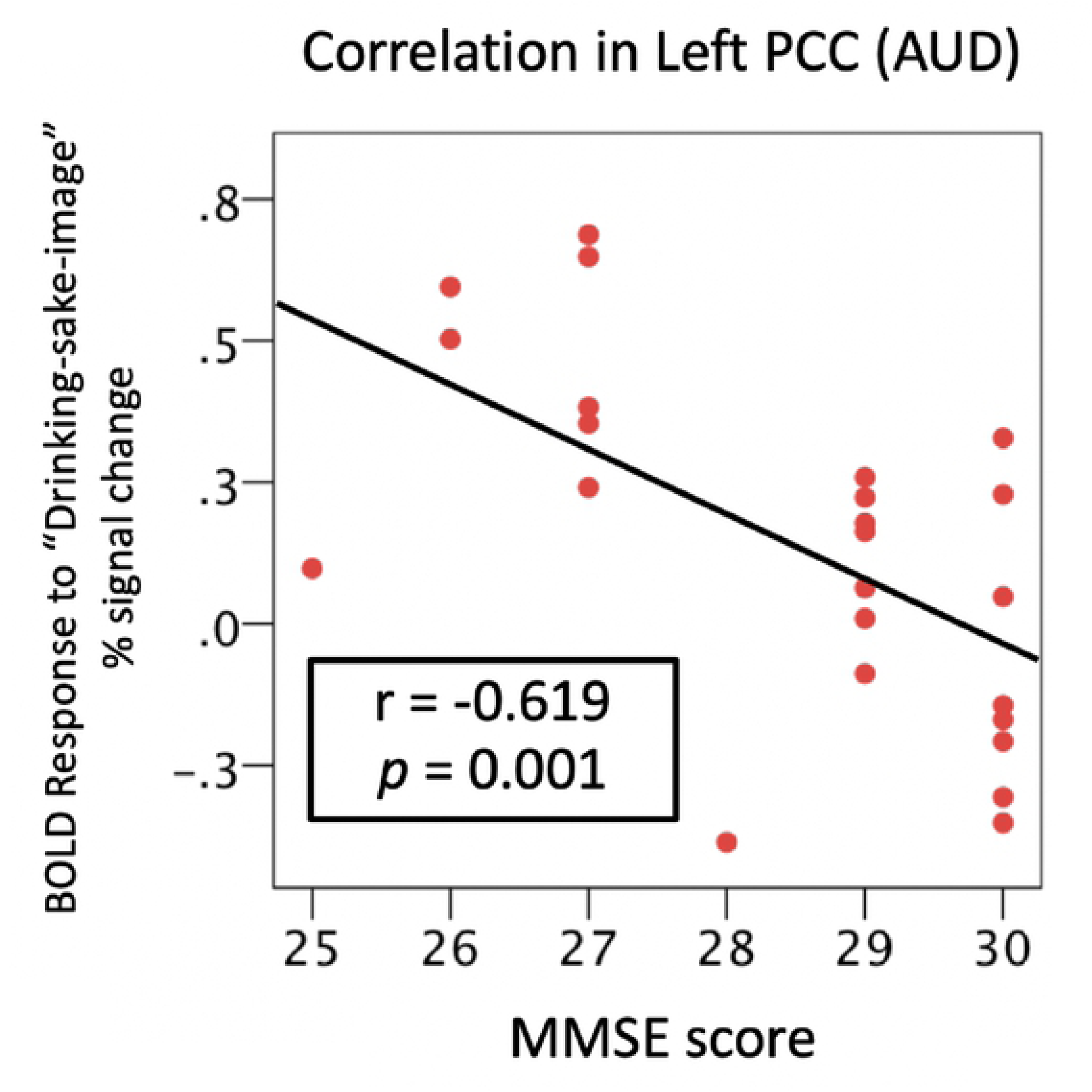
Correlation between the Mini–Mental State Examination (MMSE) scores and BOLD responses to drinking sake in AUD participants in the left PCC. When we used Spearman correlations, we found a significant negative correlation between BOLD responses and MMSE scores in the left PCC (*r* = −0.619, *p* = 0.001). We also plotted a least squares line for the convenience of the reader.

### Correlations between brain activation and demographic/clinical measurements

In the AUD participants, we found no statistically significant between-group differences in the correlations between brain activation elicited by beverage images and demographic measurements (age, sex, handedness, and education) or clinical measurements (duration of illness, number of hospital admissions, past history of neurologic symptoms, preferred alcoholic beverage, and dose of antipsychotics, acamprosate, and antidepressant drugs). In the HC participants, we also found no statistically significant between-group differences in the correlations between brain activation and demographic measurements.

## Discussion

We collected fMRI data for admitted AUD patients and HCs while they viewed substantial and behavioural visual cues of alcohol and non-alcohol beverages. To the best of our knowledge, this is the first study to demonstrate biological differences in brain function associated with strong visual behavioural cues regarding alcohol and non-alcohol beverages in AUD patients. As we expected, the AUD patient group had greater activation in the PCC to the drinking alcohol stimulus. This suggests that patients with AUD had stronger responses to the stimulus associated with alcohol drinking behavior than to a stimulus that simply contained an image of an alcoholic beverage.

In our study, the patients with AUD had lower BOLD activation with behavioural cue reactivity to the non-alcoholic beverages in the left PCC and the left precuneus. A recent meta-analysis of fMRI studies using alcohol cue reactivity demonstrated BOLD activation in several brain regions in AUD patients [19,20], including the PCC and precuneus. This suggested that the PCC plays an important role in addiction and relapse [21]. When the PCC is activated by a visual stimulus, we extract information from our episodic memory and integrate this with existing knowledge. The human precuneus is associated with several basic cognitive activities in the resting-state condition. These include the collection and evaluation of information, self-referential mental activity, extraction of episodic memory, emotion, and anxiety [22]. Previous studies have found AUD patients to have impairments in the PCC and precuneus. Our study suggests that in patients with AUD, the functional changes in the PCC may affect the circuits involved in episodic memory.

In the left precuneus, we found that the time course of brain activity in the AUD and HC groups significantly differed with respect to the behavioural cue reactivity to drinking juice, especially during the period from 15 s to 18 s and from 18 s to 21 s after the onset of the drinking juice stimulus presentation. These results indicated that in the left precuneus, the patients with AUD experienced deactivation in the latter half of the drinking non-alcohol trials. Additionally, these data suggested that the patients exhibited decreased responses in the precuneus when they viewed an unfavourable stimulus, leading to a delayed response. The precuneus is associated with the evaluation of information and emotion. The AUD group had lower BOLD responses with behavioural cue reactivity to the non-alcohol image compared with the HC group. Thus, the precuneus of the patients with AUD may have only weakly evaluated the drinking non-alcohol stimulus.

The BOLD responses in the left PCC to the alcohol-drinking stimulus were negatively correlated with MMSE scores among the patients with AUD. Considering the negative correlation with the MMSE scores, patients with decreased cognitive function may experience stronger episodic memory when they see an alcohol-drinking stimulus. Previous studies have shown that the DMN, including the PCC and precuneus, is an important component of cognitive function. Thus, the PCC and precuneus may be associated with abnormal cognitive function in AUD patients.

In interpreting the current study, it is important to consider several possible limitations. First, as most of the patients we recruited were severe AUD patients (23 severe and one moderate according to the DSM-5 criteria), selection bias should be considered. Mild and moderate AUD patients should be examined using the same approach to determine whether the BOLD activity that we observed was a state-dependent endophenotype. Second, given that the sample size of participants was small, the statistical power for assessing brain activity was limited. Future studies with larger numbers of participants are needed to determine whether brain BOLD activity can differentiate BOLD patterns elicited by the behavioural cue reactivity towards alcohol and non-alcohol beverages in AUD patients. Third, the choice of the ROIs in the functional MRI could be considered to be arbitrary. The regions were selected on the basis of their involvement in specific cue-elicited BOLD responses. The choice of ROIs could have been broadened to include other regions known to be involved in reward circuitry associated with addiction (e.g., ACC, ventral striatum, and amygdala). Despite these limitations, this line of research may contribute to the understanding of the neural basis of the addiction circuit, which may have important implications for our understanding of alcohol addiction pathology and treatment.

## Conclusions

Our study showed that, compared with controls, patients with severe AUD exhibited signs of functional brain impairment. Taking a harm reduction approach, the APA and domestic guidelines in Japan suggest that not only abstinence from alcohol use but also reduction or moderation of alcohol use may be appropriate initial goals for AUD treatment. Clinically, individuals with severe AUD generally have great difficulty moderating alcohol use. This may be related to the neural differences indicated by our study. Even clinicians who are able to form strong relationships with patients may face challenges in helping those who want to drink alcohol in moderation. Our results may help to explain the challenges encountered in such moderation efforts.

## Author contributions

S.F., H.K., N.O., T.Y., H.O., and T.U. designed the study; S.F., H.K., N.O., and T.U. performed the research; S.F., H.K., N.O., and T.U. analysed the data; S.F., H.K., N.O., and T.U. wrote the paper; S.F., T.M., and T.Y. recruited participants and acted as clinicians.

## Acknowledgments

The authors would like to thank all participants who took part in our study, as well as N. Nakayama, S. Morita, K. Kawakami, K. Tashiro, T. Yoshimori, and K. Endou for their support as research assistants. We thank The AJE Team (www.aje.com) for editing a draft of this manuscript.

